# The role of cholinergic signaling in multi-sensory gamma stimulation induced perivascular clearance of amyloid

**DOI:** 10.1101/2024.11.27.625739

**Authors:** Nicolas Lavoie, Cristina Blanco-Duque, Martin Kahn, Hiba Nawaid, Anjanet Loon, Alexander Seguin, Ravikiran Raju, Alexis Davison, Cheng-yi Yang, Li-Huei Tsai

**Affiliations:** Picower Institute for Learning and Memory, Massachusetts Institute of Technology, Cambridge, MA 02139, USA; Department of Brain and Cognitive Sciences, Massachusetts Institute of Technology, Cambridge, MA 02139, USA; Department of Pediatrics, Massachusetts General Hospital, Boston, MA 02114, USA

**Author notes:** These authors contributed equally to this work (Nicolas Lavoie, Cristina Blanco-Duque, Martin Kahn).

## Abstract

Modulatory neurotransmitters exert powerful control over neurons and the brain vasculature. Gamma Entrainment Using Sensory Stimuli (GENUS) promotes amyloid clearance via increased perivascular cerebral spinal fluid (CSF) flux in mouse models of Alzheimer’s Disease. Here we use whole-brain activity mapping to identify the cholinergic basal forebrain as a key region responding to GENUS. In line with this, GENUS promoted cortical acetylcholine release, vascular dilation, vasomotion and perivascular clearance. Inhibiting cholinergic signaling abolished the effects of GENUS, including the promotion of arterial pulsatility, periarterial CSF influx, and the reduction of cortical amyloid levels. Our findings establish cholinergic signaling as an essential component of the brain’s ability to promote perivascular amyloid clearance via non-invasive sensory stimulation.

## INTRODUCTION

Clearance of toxic molecules via perivascular spaces is essential for maintaining brain homeostasis and counteracting accumulations of toxic molecules, such as amyloid ^1–5^. Indeed, Alzheimer’s disease (AD) is associated with reduced perivascular clearance and experimental disruption of perivascular clearance leads to accumulation of amyloid and aggravates cognitive impairments ^6–8^.

GENUS has been shown to reduce amyloid pathology and cognitive symptoms in AD models across a multitude of studies from multiple independent laboratories ^9–16^. Two recent studies showed that GENUS reduces amyloid by promoting perivascular clearance. Specifically, GENUS promotes arterial pulsatility, which leads to increased cerebral spinal fluid (CSF) flux ^17,18^.

Previous studies on the mechanism of GENUS have focused on local signaling molecules, such as VIP and adenosine ^17–19^, which are produced locally in response to local neuronal responses. However, in addition to local neuronal activity, sensory processing is highly parallelized, and GENUS could activate an array of subcortical neuromodulatory regions, such as the cholinergic basal forebrain.

Subcortical neuromodulators profoundly modulate neuronal activity and the brain vasculature ^20^. For instance, acetylcholine can promote vasodilation by acting directly on endothelial cells or smooth muscle cells surrounding arteries. Interestingly, changes in vessel diameter induced by sensory stimulation were recently shown to increase arterial pulsatility and periarterial CSF flow^21^. However, a direct relationship between acetylcholine dynamics and perivascular clearance has gone unexplored.

We hypothesized that GENUS could activate extracortical systems that will contribute towards the beneficial cortical effects reported previously.

## RESULTS

### The cholinergic basal forebrain is activated in response to GENUS

We first established a brain-wide activity map to unbiasedly identify brain regions that may contribute towards the effects of GENUS on glymphatic flux and amyloid clearance. To this end we employed whole-brain tissue clearing with SHIELD ^22^ combined with brain-wide cFos labeling using eFLASH ^23^ (Fig. 1A) in 5xFAD mice, where 1 h of GENUS is known to reduce amyloid. We acquired whole-brain images via light-sheet microscopy, segmented cFos-positive cells using a 3D U-Net-based deep learning algorithm ^24^, and registered cell coordinates to the Allen Brain Atlas (Fig 1B) ^25^.

**Figure 1.**
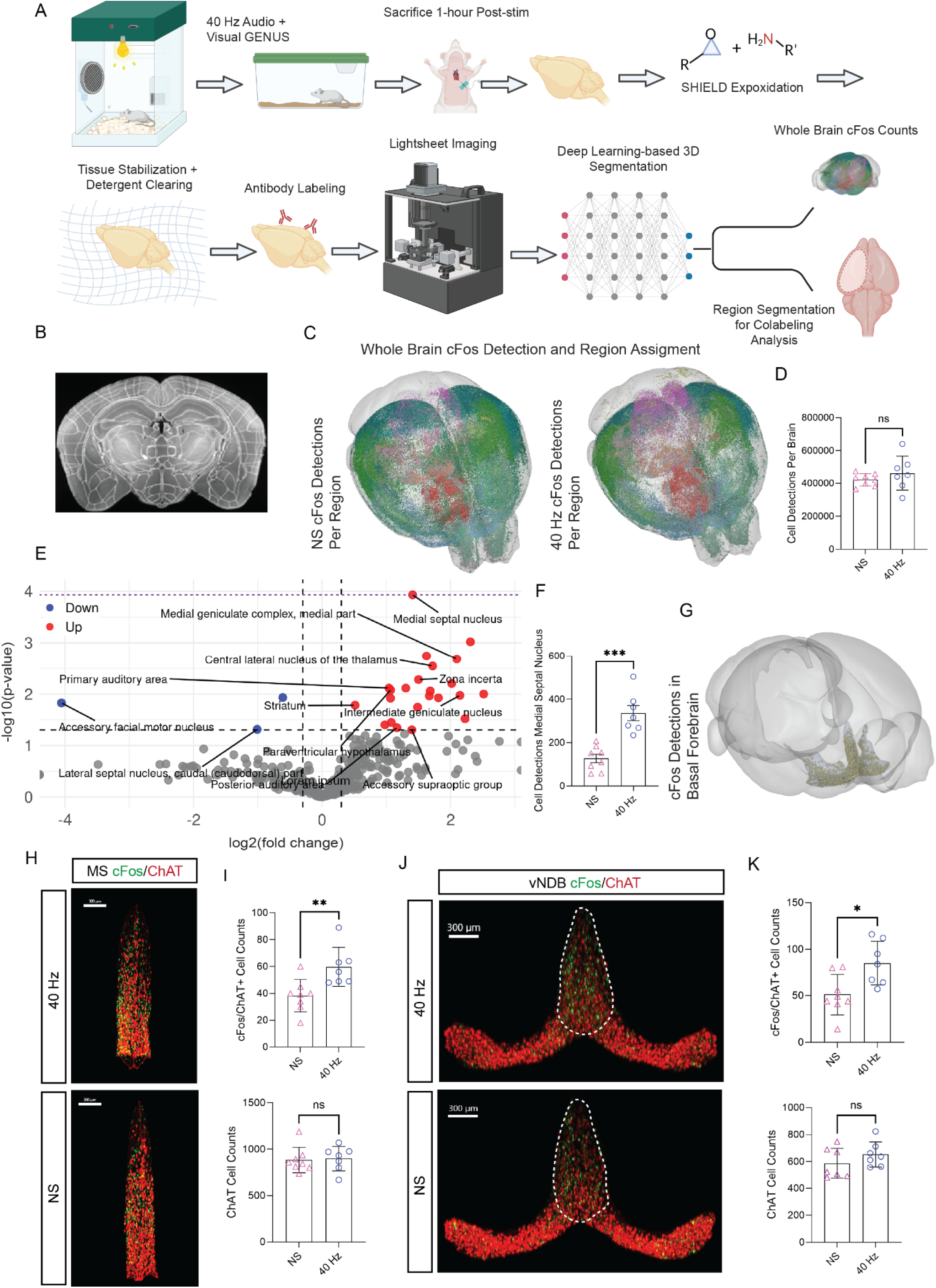
GENUS activates specific neural populations and increases cholinergic neuron activity in the basal forebrain. **(A)** 5xFAD mice were exposed to GENUS audio and visual stimulation or no stimulation, followed by sacrifice 1-hour post-stimulation. Tissue was stabilized and cleared via SHIELD tissue clearing, labeled with antibodies by eFLASH active labeling, and imaged via lightsheet microscopy. Deep learning-based 3D segmentation was employed to quantify whole-brain cFos counts, after which cFos detections were assigned to brain regions in the Allen Brain Atlas, **(B)** Representative coronal section showing Allen Brain Atlas segmentations mapped onto an acquired brain scan, **(C)** Representative whole-brain cFos detection and regional assignment in mice with and without GENUS. cFos detections were visualized as color-coded dots corresponding to their assignment in the Allen Brain Atlas, **(D)** Quantification of total cFos detections per brain in GENUS (n=7) versus no stimulation (n=8) conditions shows no significant difference (NS, unpaired t-test), **(E)** Volcano plot showing changes in cFos expression across brain regions. GENUS increased cFos levels significantly in regions including the medial septal nucleus, medial geniculate complex, and suprageniculate nucleus (red points, p < 0.05), **(F)** cFos detections in the medial septal nucleus were significantly increased in GENUS stimulated mice compared to controls (*** p < 0.0001, unpaired t-test, FDR < 0.05), **(G)** 3D rendering showing cFos detections in the basal forebrain, **(H)** Whole-mount medial septal nucleus (MS) segmentations stained for cFos (green) and ChAT (red) in GENUS and no-stimulation conditions, **(I)** Quantification of cFos+/ChAT+ double-positive cell counts in the MS revealed a significant increase with GENUS (** p < 0.01, unpaired t-test), whereas total ChAT+ cell counts were unaffected (ns), **(J)** Whole-mount vNDB segmentations showing increased cFos+/ChAT+ colocalization in GENUS stimulated mice compared to controls, **(K)** Quantification of cFos+/ChAT+ cell counts (top) and total ChAT+ cell counts (bottom) in the vNDB confirms GENUS-induced activation of cholinergic neurons (* p < 0.05, unpaired t-test, ns = not significant). Scale bars: 300 *µm*. Error bars represent mean ± SEM.

Our analysis revealed that GENUS does not produce significant overall increases in cFos positive cells relative to the no-stimulation control condition (Fig. 1C, D). Region-specific mappings, however, showed that GENUS significantly increased activity (Fig. 1D red points) in areas associated with auditory and visual processing, such as the Primary Auditory Area, Posterior Auditory Area, Medial Geniculate Nucleus, Lateral Geniculate Nucleus, and others (Fig. 1D, Supplemental Fig. 1A). Strikingly, the statistically most significant increase in neuronal activity occurred in the Medial Septal Nucleus of the Basal Forebrain (Fig. 1E, F).

To validate our whole brain cFos experiment, we co-stained brains for cFos and choline acetyltransferase (ChAT), a marker for cholinergic neurons, to confirm that GENUS activated Basal Forebrain cholinergic neurons. After aligning image stacks to the Allen Brain Atlas, we segmented Basal Forebrain regions, including the Medial Septal Nucleus, Diagonal Band Nucleus, and Substantia Innominata, and analyzed cFos/ChAT co-positive cells (Fig. 1G). We further subdivided the Diagonal Band Nucleus into the vertical and horizontal bands to provide a more granular representation of ChAT activity. Recent reports suggest that these regions exhibit distinct projection patterns, with the vertical band predominantly targeting cortical layers I-III, while the horizontal band projects to deeper cortical layers and the olfactory bulb ^26^.

We observed a significant increase in cFos/ChAT colocalization in the Medial Septal Nucleus of mice treated with GENUS, without a significant change in the overall number of ChAT+ cells (Fig. 1H, I), consistent with the whole-brain cFos data. Similarly, increased cFos/ChAT+ cells were detected in the vertical Diagonal Band Nucleus, again with no change in ChAT+ cell numbers (Fig. 1J, K). However, no significant increase in cFos/ChAT+ cells was observed in the horizontal Diagonal Band Nucleus or Substantia Innominata (Supplementary Fig. 1B, C).

### GENUS promotes acetylcholine release in frontal cortex

The vertical Diagonal Band Nucleus (vDBN) sends cholinergic axons to frontal cortex, where we previously observed increased perivascular clearance in response to GENUS (ref). Therefore, we next tested if GENUS leads to acetylcholine release using fiber photometry recordings of the GRABAch3.0 ^27^ sensor (Fig. 2A). We observed a significant increase in acetylcholine release compared to baseline. This effect was abolished when mice were injected with the muscarinic receptor blocker scopolamine, which directly blocks the sensor and thus, confirms that the response to GENUS indeed represents cholinergic signaling and not changes in background fluorescence. (Fig. 2B, 2C).

**Figure 2.**
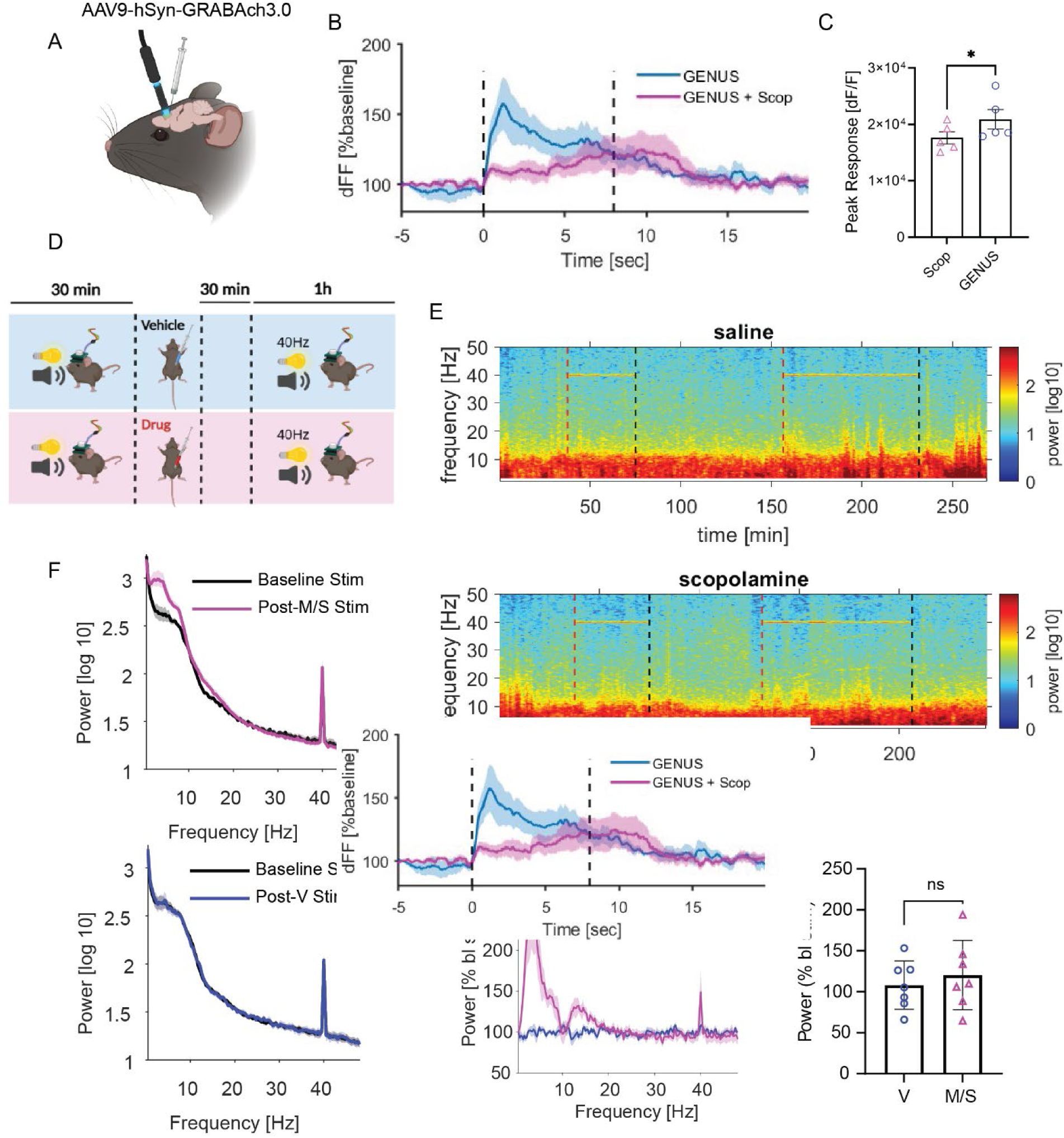
GENUS induces acetylcholine release in the frontal cortex, but acetylcholine blockade does not disrupt entrainment. **(A)** Schematic of AAV9-hSyn-GRAB-ACh3.0 injection into the frontal cortex to measure acetylcholine dynamics via fiber photometry during GENUS, **(B)** Fiber photometry traces showing the change in fluorescence (ΔF/F, %) in response to GENUS with and without scopolamine (Scop). Scopolamine attenuates the acetylcholine response to GENUS, **(C)** Quantification of peak acetylcholine responses to GENUS with and without scopolamine (*p < 0.05, paired t-test), **(D)** Experimental design to evaluate the effects of saline (vehicle) or the cholinergic antagonists scopolamine and mecamylamine (M/S) on GENUS. Mice were treated with vehicle or M/S 30 minutes prior to stimulation, **(E)** Spectrograms showing power density of neural activity across frequencies during GENUS in vehicle or M/S-treated conditions. Both saline and M/S-treated mice exhibit robust 40 Hz power during stimulation, **(F)** Power spectra comparing baseline stimulation to post-stimulation in mice treated with vehicle (top) or scopolamine (bottom). GENUS-induced 40 Hz power is unaffected by injection of cholinergic antagonists, **(G)** Normalized power spectra during GENUS in vehicle and M/S-treated mice, showing no significant difference in power at 40 Hz in M/S-treated mice, **(H)** Quantification of 40 Hz power during GENUS in vehicle versus M/S-treated mice. No significant differences were observed between groups (* p > 0.05, unpaired t-test. Error bars represent mean ± SEM.

Cholinergic signaling can be involved in the induction and maintenance of endogenous gamma oscillations ^28^. However, it is not clear to what extent gamma-range neuronal activity evoked by GENUS mechanistically overlaps with endogenous gamma oscillations ^29^. To assess the potential contribution of cholinergic signaling to GENUS-induced neuronal activity, we combined *in vivo* electrophysiology recordings with pharmacological inhibition of cholinergic signaling. We recorded cortical EEG in freely moving 5xFAD mice for 30 min and then administered saline or a combination of cholinergic antagonists (scopolamine and mecamylamine). Mice were then exposed to one hour of GENUS (Fig. 2D). As expected from prior work, cholinergic antagonists increased the power of slower frequencies, which is consistent with the desynchronizing effect of cholinergic signaling. However, we did not observe a difference in the stimulation-induced 40 Hz power band (Fig. 2E-H).

### Cholinergic signaling contributes to CSF influx induced by GENUS

Previous research has demonstrated that GENUS facilitates perivascular clearance by increasing influx of CSF influx and arterial pulsatility^17^. To assess whether cholinergic activity is involved in these processes, we investigated the effects of GENUS on CSF influx and arterial pulsatility in mice treated with the cholinergic antagonists scopolamine and mecamylamine.

To assess CSF influx, we injected fluorescent tracer (dextran, 3 kDa) into the cisterna magna of 5XFAD mice and monitored tracer influx into frontal cortex using two-photon microscopy. We treated mice with cholinergic antagonists 30 minutes before monitoring CSF influx during one hour of GENUS (Fig. 3A). Consistent with previous findings ^17,18^, saline-treated mice exhibited significantly greater CSF tracer accumulation in the cortex after GENUS compared to no stimulation (Fig. 3B, C). However, this increase was absent in mice treated with cholinergic antagonists, indicating that cholinergic signaling is required for stimulation-induced perivascular influx into the brain parenchyma (Fig. 3B, C).

**Figure 3.**
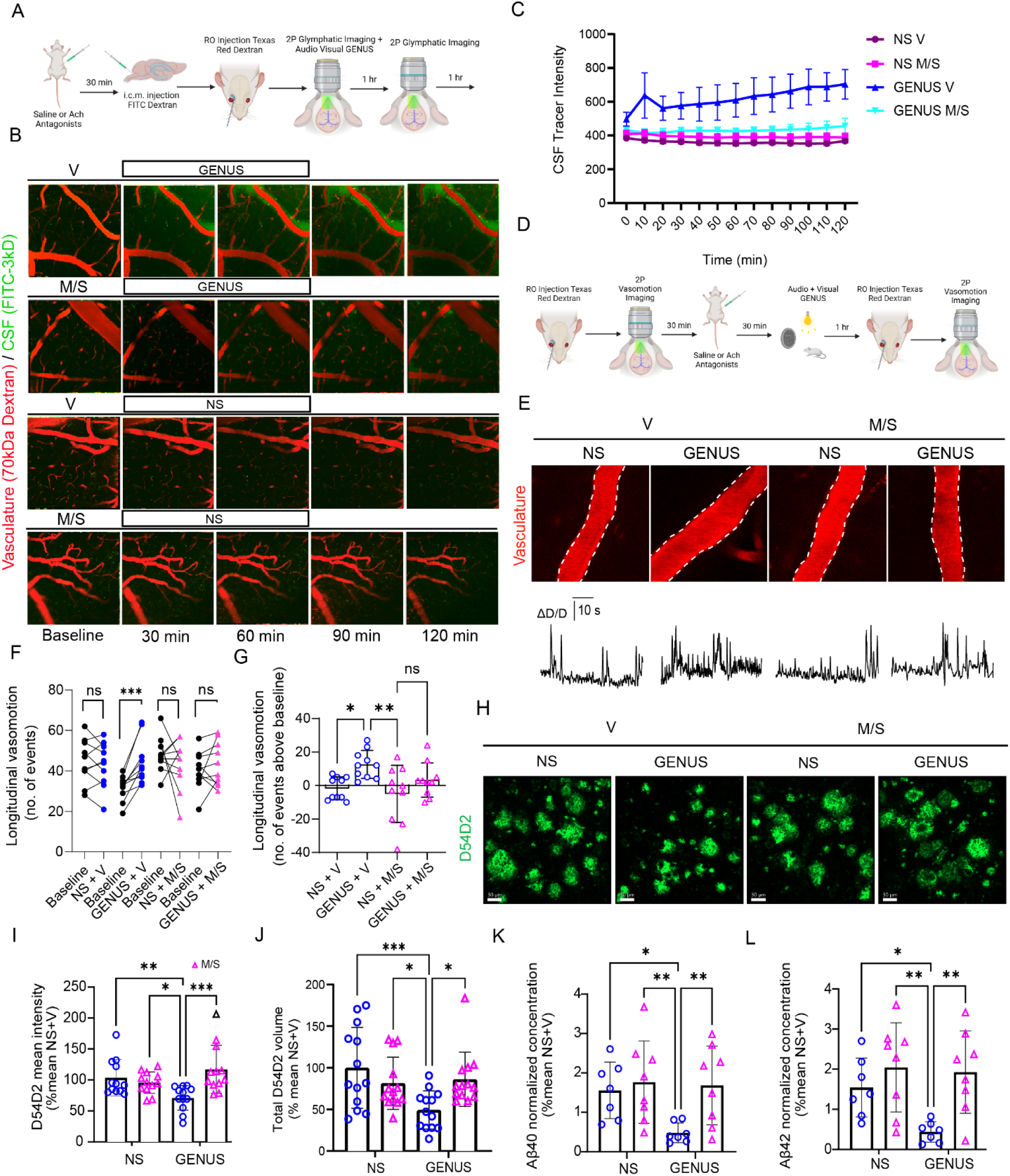
GENUS enhances glymphatic influx, vasomotion, and amyloid clearance, effects that are blocked by cholinergic antagonists. **(A)** Vehicle (saline, or V) or acetylcholine antagonists (mecamylamine and scopolamine, or M/S) were administered 30 minutes prior to cisterna magna injection of FITC-dextran tracer and retro-orbital injection of the vascular tracer, 70 kDa Texas Red dextran. Two-photon imaging was performed every 10 minutes during and after GENUS or no stimulation to measure CSF tracer influx, **(B)** Workflow for assessing vasomotion and vasodilation: Texas Red dextran was injected into the retro-orbital sinus, followed by two-photon imaging of vasculature during GENUS or no stimulation with or without acetylcholine antagonists, **(C)** Representative images of glymphatic influx (CSF tracer, green) and vasculature (Texas Red dextran, red) during GENUS or no-stimulation (NS) conditions with vehicle or M/S CSF influx is visibly enhanced during GENUS in vehicle-treated mice but is blocked by M/S, **(D)** Quantification of CSF tracer intensity over time shows significant GENUS-induced increases in vehicle-treated mice, which are suppressed by M/S (two-way ANOVA), **(E)** Representative two-photon images and traces showing changes in vascular diameter (ΔD/D) during GENUS with vehicle or M/S treatment. GENUS-induced increases in vasodilation are attenuated by M/S, **(F-G)** Quantification of longitudinal vasomotion. **(F)** No significant differences (ns) in vasomotion frequency across conditions. **(G)** GENUS significantly increases vasomotion amplitude in vehicle-treated mice (** p < 0.01), but not in M/S-treated mice (ns, one-way ANOVA with Tukey’s multiple comparison test), **(H)** Representative images of brain sections showing D54D2 amyloid tracer in GENUS and no-stimulation conditions with vehicle or M/S GENUS-induced reductions in amyloid levels are attenuated by M/S, **(I-J)** Quantification of D54D2 mean intensity **(I)** and total volume **(J)** shows significant reductions in amyloid burden during GENUS with vehicle (** p < 0.01), but not with M/S treatment, **(K-L)** Normalized ELISA quantification of Aβ40 **(K)** and Aβ42 **(L)** concentrations shows significant reductions during GENUS (** p < 0.01) with vehicle, but not with M/S. treatment. Scale bars: 30 *µm*. Error bars represent mean ± SEM.

### Cholinergic signaling is required for arterial pulsatility and vasodilation induced by GENUS

GENUS has been shown to promote arterial dilation and pulsation ^12,17,18^, processes hypothesized to facilitate CSF movement. Since acetylcholine is critical for arterial vasodilation and vasomotion, we hypothesized that blocking cholinergic neurotransmission would attenuate these vascular effects. To test this, we labeled blood vessels with Texas Red Dextran in 5xFAD mice and imaged vascular dynamics through cranial windows over the frontal cortex. Arterial vasomotion was measured for five minutes before and after mice were treated with cholinergic antagonists or saline and exposed to one hour of GENUS or no stimulation (Fig. 3D).

In agreement with prior work ^12,17,18^, saline-treated mice exhibited increased arterial diameter and pulsatility following GENUS compared to no stimulation (Supplementary Fig. 2A-C). However, this effect was abolished in mice treated with cholinergic antagonists, where arterial dynamics did not differ between stimulation and control conditions (Supplementary Fig. 2A-C). These results suggest that cholinergic neurotransmission is necessary for gamma-related arterial vasodilation and pulsation. The primary mechanism likely involves acetylcholine binding to muscarinic receptors on endothelial cells, which stimulates endothelial nitric oxide synthase (eNOS) activity and nitric oxide release, leading to vascular smooth muscle relaxation ^20^. To investigate whether GENUS enhances eNOS expression, we exposed 6-month-old wild-type mice to either one hour of GENUS or no stimulation. Mice were sacrificed immediately afterward, and Nos3 (encoding eNOS) expression in prefrontal cortex endothelial cells was quantified via in situ hybridization (Supplementary Fig. 3A). We found a significant increase in Nos3-positive puncta in GENUS-treated mice compared to controls (Supplementary Fig. 3B). Importantly, there was no difference in PECAM1-positive puncta, indicating that increased Nos3 expression was not due to differential vessel sampling. These findings demonstrate that GENUS stimulates eNOS expression, likely contributing to its effects on arterial vasodilation and CSF influx.

### Cholinergic activity is required for GENUS-mediated amyloid clearance

Given that the cholinergic system is necessary for GENUS-mediated glymphatic perivascular flux, we next asked whether it is also required for removal of parenchymal amyloid. To test this, we injected the fluorescent tracer A**β**42-HiLyte into the frontal cortex of wild-type mice ^30^. Mice were treated with either cholinergic antagonists or saline, exposed to GENUS or no stimulation and brain tissue was collected. Confocal imaging of coronal sections revealed significantly lower A**β**42 tracer levels in the parenchyma of saline-treated mice after GENUS compared to no stimulation (Supplementary Fig. 4A, B). In contrast, no difference in tracer retention was observed between stimulated and unstimulated conditions in the cholinergic antagonist group (Supplementary Fig. 4C). This suggests that cholinergic signaling is required for the clearance of exogenous amyloid.

To investigate whether cholinergic signaling is required for the removal of endogenously produced amyloid, we treated 5XFAD mice with either cholinergic antagonists or saline, exposed them to one hour of GENUS or no stimulation, and measured amyloid levels in the frontal cortex. Immunohistochemical analyses using the amyloid-specific antibody D54D2 (Fig. 3H) revealed significant reductions in both the total area covered by amyloid (Fig. 3I) and mean intensity (Fig. 3J) in mice exposed to GENUS and saline. These effects were abolished in mice that received either no stimulation or cholinergic antagonists. To corroborate these findings, we performed enzyme-linked immunosorbent assays (ELISA) for Aβ40 and Aβ42 in the frontal cortex, which similarly showed reduced amyloid levels only in mice treated with GENUS and saline (Fig. 3K, L).

## DISCUSSION

Here we show that GENUS activates cholinergic neurons in the basal forebrain and promotes the release of acetylcholine in frontal cortex. Additionally, we show cholinergic neurotransmission in frontal cortex is necessary for the ability of sensory stimulation to increase arterial pulsatility, CSF influx and the clearance of cortical amyloid.

We present the first brain-wide map of cFos activity following GENUS, which suggested GENUS differentially induces activation of the basal forebrain, which contains cortically-projecting cholinergic neurons. Furthermore, co-staining of cFos with a cholinergic marker confirmed that cholinergic neurons, which are known to project to frontal cortex and hippocampus ^26^, are activated by GENUS. Importantly, we were able to confirm the cFos-based activity mapping by means of fiber photometry recordings of cholinergic release in frontal cortex. Importantly, frontal cortex is where we previously observed the effects of GENUS on CSF flux ^17^.

Besides the activation of the cholinergic forebrain, this rich dataset revealed multiple regions that could be relevant to how GENUS propagates throughout the brain and exerts its beneficial effects. Besides the expected activation of the primary sensory processing, (e.g. medial/lateral geniculate body) other noteworthy regions include the pretectal nucleus, the paraventricular hypothalamus and the striatum. The tectum has been highlighted by a prior cFos experiment using GENUS ^16^, is a region that receives extensive unfiltered multisensory stimulation and has extensive projections. It could present a propagation route by which GENUS reaches other higher-order brain regions. The activation of the striatum is of relevance for potential applications of GENUS in Parkinson’s Disease.

Surprisingly, the role of cholinergic signaling in perivascular clearance has not been extensively investigated. It has long been known that acetylcholine has strongly vasoactive properties ^31^ and that stimulation of the basal forebrain leads to vasodilation that is mediated by cholinergic signaling ^32^. Recent studies suggest that CSF influx is driven by alternating arterial dilations and contraction ^21^ and that GENUS promotes such arterial pulsatility. We find that cholinergic antagonists abolish the ability of sensory stimulation to promote arterial pulsatility and perivascular flux. This appears unrelated to changes in baseline motility, since cholinergic antagonists did not significantly reduce vasomotive events in the absence of stimulation, compared to control conditions (Fig 3 F,G note NS +V vs NS + M/S). Similarly, cholinergic antagonists in the absence of stimulation did not appear to change baseline perivascular flux. This suggests that cholinergic signaling is important specifically for the increase in arterial pulsatility induced by sensory stimulation.

There are multiple, not mutually exclusive, mechanisms that could account for our findings. The most parsimonious is that GENUS leads to acetylcholine release in frontal cortex, which acts directly or indirectly on the neurovasculature. Acetylcholine is known to be able to directly act on endothelial cells or smooth muscle cells. In endothelial cells, cholinergic signaling leads to nitric oxide mediated vasodilation ^20^. In support of this we observe upregulated eNOS signaling in brain endothelial cells following GENUS (Fig. S3). An alternative – not mutually exclusive – mechanism is that cholinergic signaling indirectly acts on the vasculature by interacting with local non-vascular cells.

How does non-local neuromodulation integrate with locally produced signals that can regulate vasomotion and CSF flux? Prior studies have shown that VIP and adenosine are both required for enhanced vasomotion and CSF influx following GENUS ^17,18^. These messengers are likely both produced in response to local neuronal activity. Interestingly, there are established direct links between cholinergic signaling and both VIP and adenosine signaling. First, cholinergic signaling can activate VIP interneurons, which could be required for VIP mediated vasodilation described by Murdock et al ^17^. Second, adenosine is a powerful regulator of cholinergic release at the level of basal forebrain firing rates ^33^ and at the level of presynaptic release ^34^. Third, cholinergic tone can affect local neuronal firing and excitability ^35^, which could alter the production of adenosine levels. In this model, locally released neuromodulators and external acetylcholine synergistically interact to promote vasomotion and CSF flow.

Overall, our data suggests that perivascular amyloid clearance evoked by GENUS requires the recruitment of local and external neuromodulators.

Together, these results demonstrate that cholinergic neurotransmission is indispensable for gamma-mediated increases in glymphatic flow dynamics. Cholinergic signaling supports gamma-induced arterial dilation and pulsatility, facilitating CSF influx into the brain parenchyma and amyloid clearance. Gamma multisensory stimulation enhances eNOS expression, linking acetylcholine signaling to vascular and glymphatic processes. These findings highlight the critical role of cholinergic activity in mediating the therapeutic effects of gamma stimulation.

## METHODS

### Mice

Experiments were performed in 5XFAD mice (male, 6 to 11-month-old), which were bred and maintained in the MIT animal facility, and C57BL/6J mice (male, 6 to 10-month-old) obtained from Jackson Laboratory. All animal experiments adhered to NIH guidelines and were supervised by the Massachusetts Institute of Technology Institutional Animal Care and Use Committee. Mice were housed in groups of no more than five, except when they were implanted with probes for electrophysiological recordings or implanted with headbars for two-photon imaging and were single housed. The housing conditions of all mice followed the temperature (18–26°C) and humidity (30–70%) ranges outlined in the ILAR Guide for the Care and Use of Laboratory Animals (1996). Every effort was made to minimize animal usage.

### Non-invasive multisensory sensory stimulation

Audio-visual 40Hz stimulation was performed in plexi-glass cages (30 x 25 x 25cm) that were surrounded on all four sides by a LED strip programmed to emit light flickering at either 40 Hz (12.5 ms light on, 12.5 ms light off) or constant light. The light brightness within cage varied between 600 lux (middle) and 1000 lux (walls). Above the chambers, speakers (AYL, AC-48073) were positioned and programmed to produce a 10 kHz tone, delivered at 70 decibels, and with a duration of 1 ms with inter-stimulus-intervals of 24 ms to create 40Hz sound trains. Both the LED and speakers were coordinated through a microcontroller (Teensy) to ensure simultaneous delivery of sensory input, aligning stimulus pulses of each modality with the onset of each pulse. To perform the audio-visual stimulation procedure, mice were individually housed in the plexi-glass cages and habituated to the chamber for 48 hours. The stimulation was always presented during the light period starting between zeitgeber time (ZT) 2 and 5. At the time of stimulation, mice were exposed to GENUS or constant light for one hour during wakefulness. To ensure continuous wakefulness during the experiment, nestlet was removed and small novel objects were provided. This is a standard method to ensure wakefulness while minimizing stress.

### SHIELD processing and whole brain immunolabeling

Mice were sacrificed for SHIELD processing 1 hour after GENUS. To minimize circadian effects and variability in post-stimulation peak cFos expression ^36–39^, mice were grouped so that n=2 (n=1 per group) would undergo stimulation per day at ZT 3. For SHIELD tissue processing ^22,40^, mice were perfused with ice-cold heparinized (20 U/mL, Sigma) 1× PBS followed by ice-cold 4% paraformaldehyde (w/v). Brains were carefully dissected to avoid tissue damage. Samples were then processed according to the manufacturer’s instructions (LifeCanvas Technologies, USA). Briefly, they were placed in SHIELD OFF at 4°C for 3 days, followed by SHIELD ON at 37°C for 24 hours. Samples were actively cleared with Delipidation Buffer for 30 hours in a SmartBatch (LifeCanvas Technologies, USA). After delipidation, the tissue was blocked with Blocking Solution (LifeCanvas Technologies, USA) for 48 hours at 37°C and actively labeled with cFos (Abcam, ab214672, UK) and ChAT (Millipore, AB144P, USA) antibodies. Finally, samples were refractive index-matched with EasyIndex (LifeCanvas Technologies, USA), embedded in agarose, and imaged using a SmartSPIM lightsheet microscope (LifeCanvas Technologies, USA) with a 3.6x objective at a resolution of 1.8 µm/pixel in XY and 4 µm in Z throughout the brain.

### Quantitative cFos activity mapping

Raw lightsheet images were destriped using pystripe (v 1.20) and stitched with Terastitcher ^41,42^. Whole brain datasets were then registered to the Allen Brain Atlas using the autofluorescence channel. This was done by first applying CLAHE adaptive histogram equalization, followed by registration with the python package brainreg (v 1.0.10) ^25,43,44^, which enables accurate, fully automated Allen Brain Atlas registration. To quantify brainwide cFos counts, we employed cellfinder (v 1.3.1) ^24^, which uses a 3D U-Net for cell segmentation and a ResNet architecture to classify segmentations as either cFos-positive or -negative. This method, optimized for 3D segmentation in terabyte-sized datasets, was validated in 5 patches (100x100x100), showing 95% accuracy in classifying cFos-positive cells. Cell detections were visualized with brainrender (v 2.1.10) ^43^, and custom python scripts were developed to pseudocolor detected cells based on their assigned Allen Atlas region and to extract detections assigned to the basal forebrain. Additionally, we wrote custom python scripts to extract brain region segmentations and quantified cFos/ChAT colocalization by creating cell surfaces in Imaris (v 10.2, Bitplane, Oxford Instruments, Switzerland).

### Pharmacology

To modulate cholinergic activity in mice, we used the nicotinic receptor antagonist mecamylamine (3mg/kg) and the muscarinic receptor antagonist scopolamine (1mg/kg) administered via an intraperitoneal injection 30 minutes prior to sensory stimulation. We used this dose based on prior literature of acetylcholine activity modulation ^45,46^.

### EEG and EMG surgical implantation

Surgeries were performed under anesthesia with isoflurane (induction, 5%; maintenance, 2%). Analgesics, including Metacam (5 mg/kg) and Buprenorphine (1 mg/kg), were administered and ophthalmic ointment (Puralube Vet Ointment, Dechra) was applied to safeguard the cornea. Animals were secured in a stereotactic frame (Kopf Instruments) with a heating pad maintaining body temperature at 37°C. The scalp fur was shaved, and the area was cleaned with betadine and 70% ethanol. Lidocaine (0.05 mL, 5 mg/ml) was topically applied as a local analgesic, and a surgical scalpel (Fine Science Tools) was used to make an incision across the scalp (AP) and on the skin overlaying the nuchal muscle. A high-speed hand dental drill (0.7mm burr, Fine Science Tools) was used to make skull holes above frontal (+2mm AP, +2mm ML, relative to bregma), somatosensory (−1mm AP, +2mm ML), occipital (−4mm AP, +2.5-mm ML) and cerebellar (−7mm AP, +1mm ML) brain regions. The EEG and electromyography (EMG) recording mounts were then implanted. These mounts consisted of stainless-steel screws (shaft diameter 0.86mm, InterFocus Ltd, UK) and two single-stranded stainless-steel wires connected to an 8-pin mount connector (8415-SM, Pinnacle Technology Inc, USA), as detailed previously ^47,48^. Epidural screws were secured into the holes for EEG signal recordings and tungsten wires were inserted in the nuchal muscle to record the EMG. A reference screw was implanted over the cerebellum. All screws and wires were affixed to the skull using dental cement (C&B Metabond, Parkell Inc).

### EEG and EMG data acquisition

To conduct electrophysiology recordings, mice were individually housed in plexi-glass cages (30 x 25 x 25cm) situated within sound-attenuated and ventilated Faraday chambers (Med Associates, Inc). For acclimatization, mice were placed in these chambers at least 48h before experiments. During this time, animals were habituated to tethered recording conditions. Electrophysiological recordings were acquired with a RZ2 High Performance Processor and Synapse software (Tucker-Davis Technologies Inc., USA). EEG and EMG signals were continuously recorded, amplified, and digitized with a Subject Interface Module (Tucker-Davis Technologies Inc., USA), and saved in a local computer. EEG and EMG signals were filtered between 0.1–100Hz and stored at a sampling rate of 305Hz. The signals were resampled offline at a sampling rate of 256 Hz using custom-made Matlab (The MathWorks Inc, Natick, Massachusetts, USA) scripts. For subsequent analyses, EEG and LFP power spectra were computed by a Fast Fourier Transform of 4-s epochs (Hanning window), with a 0.25Hz resolution Matlab (The MathWorks Inc, Natick, Massachusetts, USA).

### Cranial window implantation

Surgical procedures were performed under anesthesia with isoflurane (induction, 5%; maintenance, 2%). Analgesics, including Metacam (1 mg/kg) and Buprenorphine (0.05 mg/kg) were administered. Ophthalmic ointment (Puralube Vet Ointment, Dechra) was applied to protect the cornea. Animals were secured in a stereotactic frame (Kopf Instruments) with a heating pad to maintain body temperature at 37°C. The scalp fur was shaved, and betadine and 70% ethanol were used to clean the area. Lidocaine (0.05 mL, 5 mg/ml) was topically applied as a local analgesic and a surgical scalpel (Fine Science Tools) was used to perform a small incision on the scalp. Dental cement (C&B Metabond, Parkell Inc) was then used to attach a titanium headplate to the skull. The headplate was centered around the frontal cortex (+2mm anterior-posterior (AP) +0.3mm medio-lateral (ML), relative to bregma). A ∼4-mm circular section of the skull was removed using a drill (0.5-mm burr, Fine Science Tools). Saline was constantly flushed to keep the area clean. A 3-mm glass coverslip (Warner Instruments) was then placed over the brain, and veterinary adhesive (Vetbond, Fisher Scientific) formed a seal between the coverslip and the skull. Dental cement (C&B Metabond, Parkell Inc) was then used as a protective cover on top of the veterinary adhesive. Recovery took place on a 37°C heating pad and the analgesic Metacam (1 mg/kg) was administered 24 hours post-surgery for 3+ days. Mice had a recovery period of 4 weeks before experiments.

### Intra cisterna magna cannulation

We followed established procedures described in ^17^. The surgery was performed under anesthesia isoflurane (induction, 5%; maintenance, 2%). Opthalmic ointment and analgesics, including Metacam (1 mg/kg) and Buprenorphine (0.05 mg/kg), were administered. The mouse was secured in a stereotaxic frame (Knopf) with the head tilted to form a 120° angle with the body. The scalp and neck fur were shaved, and betadine and 70% ethanol were used to clean the area. The occipital bone was identified, and a scalpel was used to make a ∼1 cm incision on the overlaying skin. The cisterna magna was exposed by pulling apart the superficial connective tissue and neck muscles using sterile forceps. A 30G needle, prepared with PE10 tubing (Polyethylene Tubing 0.024′ OD x 0.011′ ID, BD Intramedic) and filled with fresh artificial CSF, was inserted through the dural membrane, ensuring no damage to the cerebellum and medulla. Cyanoacrylate glue (Loctite) was used to attach the cannula into the dural membrane. Dental cement (C&B Metabond, Parkell Inc) was then used to secure the needle in place. The tubing was sealed with a cauterizer (Fine Science Tools). Recovery took place on a 37°C heating pad and CSF tracer infusion, cholinergic antagonist injections, and gamma stimulation were performed following the cannulation.

### In vivo two-photon imaging

In vivo two-photon imaging was performed in awake head-fixed mice. A flat running wheel was 3D printed, covered in waterproof neoprene foam, and placed below a custom head fixation device. This device was designed using titanium bars and fork heads (Thor labs). Mice underwent a 5-day acclimatization period during which they were habituated to gentle handling, the running wheel and head-fixation. During head-fixation, mice were secured to the titanium head fork using #0-80 screws. Before imaging, the cranial window was cleaned with a cotton-tipped swab and covered with ∼1 mL dollop of Aquasonic Clear Ultrasound Transmission Gel (Parker). Two-photon microscopy images were captured using an Olympus FVMPE-RS microscope. High-resolution and high-numerical aperture imaging was utilized for data acquisition.

### Two-photon imaging of CSF tracer

Two-photon imaging of CSF was conducted on head-fixed 5XFAD and WT mice that had received cranial window implantation overlaying the frontal cortex four weeks earlier, undergone a five-day habituation period, and had intra-cisterna magna cannulation performed one hour before the imaging session. Mice were then injected with either cholinergic antagonists (scopolamine and mecamylamine) or saline, and after 30 minutes, 10 µl of a 0.5% concentration fluorescent CSF tracer (fluorescein-conjugated dextran, 3kD, Invitrogen #D3306) in artificial CSF^17^ was administered through the cannula at a rate of 1 μl/min over a period of 10 minutes^17^. The tube was then sealed with a handheld cauterizer (Fine Science Tools, #18010-00). Mice then received a retro-orbital injection of Texas Red Dextran 70kD (to label blood vessels) and were head-fixed. To visualize CSF tracer movement and blood vessels, a Spectra-Physics InsightX3 DeepSee laser tuned to 920 nm was employed. Fluorescence was collected using a 25X, 1.05 numerical aperture water immersion objective (Olympus) for one hour of either 40Hz audio-visual stimulation or no stimulation and signal detection was carried using Fluoview acquisition software (Olympus). Simultaneous image acquisition in the 575-645 nm range (for vascular structure visualization) and the 495-540 nm range (CSF tracer visualization) was conducted Z-stacks were imaged (100 µm from the cortical surface; imaging rate set to 2.0µs/pixel for the 512x512 pixel region, covering ∼509.117 µm2). Tracer influx was quantified by a blinded investigator using ImageJ, and average fluorescence intensity was calculated between z-stacks and normalized to non-treated mice.

### Two-photon imaging of arteriole pulsation

Two-photon imaging of arteriole pulsation was conducted on head-fixed 5XFAD and WT mice. These mice had undergone cranial window implantation over the frontal cortex four weeks earlier and had additionally experienced a five-day habituation period to the two-photon apparatus. 30 minutes before an imaging session, mice were either injected with saline or cholinergic antagonists (scopolamine and mecamylamine). To visualize the vasculature, a Spectra-Physics InsightX3 DeepSee laser tuned to 920 nm was utilized. Fluorescence was collected through a 25X, 1.05 numerical aperture water immersion objective (Olympus), and signal detection was performed using Fluoview acquisition software (Olympus). We recorded time-series of arterial pulsatility using a resonance scanner. In total, 5,000 frames were captured with an imaging rate of 65.779 ms/frame immediately before and after either 40Hz stimulation or no stimulation. Each recording spanned 328.90 seconds, covering an area of 160.7µm² at a rate of 0.067 ms/pixel and 0.127 ms/line. To mitigate motion artifacts, we used the phase correlation rigid registration method in suite2p ^49^ using a Gaussian smoothing of 1.15 after phase correction and 1000 frames per batch. To assess arterial pulsatility, a perpendicular section of the artery was converted to binary format using ImageJ, and the diameter segment was measured using Python. A Savitzky-Golay filter (window size 7, polynomial order 5) was employed on the vasomotion trace, and peaks were identified using a high threshold at 0.3, as a fraction of the maximum height.

### Ex-vivo fluorescence imaging of Aβ_1-42_

A fluorescent Aβ_1-42_ tracer (Beta-Amyloid (1-42), HiLyte™ Fluor 555-labeled) was prepared as 4 mg/mL solution in PBS and injected into the frontal cortex of 5XFAD mice. For the injection, mice were anesthetized with isoflurane (induction, 5%; maintenance, 2%). Analgesics, including Metacam (1 mg/kg) and Buprenorphine (0.05 mg/kg), were administered and ophthalmic ointment (Puralube Vet Ointment, Dechra) was applied to protect the cornea. Mice were secured in a stereotactic frame (Kopf Instruments) with a heating pad maintaining body temperature at 37°C. The scalp fur was shaved, and the area was cleaned with betadine and 70% ethanol. Lidocaine (0.05 mL, 5 mg/ml) was topically applied as a local analgesic, and a surgical scalpel (Fine Science Tools) was used to make an incision across the scalp (AP). A high-speed hand dental drill (0.5mm burr, Fine Science Tools) was used to make a skull hole above the frontal cortex (+2mm AP, +2mm ML, relative to bregma). A Hamilton syringe with 33-gauge needle was then lowered to 1.3 DV from the skull and after 30 seconds it was slowly retracted up to 1.2 DV were 70 nL of the A**b**_1-42_ tracer solution was injected at 15 nL/min using a syringe pump (WPI). After 2 minutes, the needle was further retracted to 1.1 DV where 70 nL were injected at 15 nL/min. Five minutes after the end of the injection, the needle was retracted to 1.0 DV and after 5 minutes the needle was slowly retracted from the brain completely. The incision was sutured, and the mice were removed from isoflurane 2 minutes after the needle retraction. Recovery took place on a 37°C heating pad. Subsequently, animals were placed in a chamber for one hour of noninvasive multisensory stimulation or control, followed by an additional 30 minutes. To prevent potential confounds associated with death by substances like isoflurane affecting glymphatic flow, mice were then euthanized through cervical dislocation. The brains were extracted and fixed overnight by immersion in 4% paraformaldehyde in PBS at 4°C. For visualizing tracer movement within the brain parenchyma, brain sections of 40 µm thickness were obtained using a vibratome (Leica). Fluorescence imaging was performed on a LSM 900 confocal microscope (Zeiss). Tracer efflux was quantified by a blinded investigator using the FIJI image processing software (v1.54, NIH). The total fluorescence intensity was computed across nine slices for an individual animal, leading to a singular biological replicate. Consistent coronal brain slices were employed for all biological replicates.

### Tissue collection and immunohistochemistry

Following the experimental procedures, mice were given a lethal dose of isoflurane and underwent transcardial perfusion with 40 mL of ice-cold PBS, followed by 40 mL of PBS with 4% paraformaldehyde (PFA; Electron Microscopy Sciences, Cat#15714-S). Subsequently, the brains were extracted and post-fixed in 4% PFA overnight at 4°C before being transferred to PBS for sectioning. Brains sliced into 40 µm sections using a vibratome stage (Leica VT1000S). Subsequently, the slices were washed with PBS and subjected to blocking with 10% normal donkey serum and 2% bovine serum albumin in PBS containing 0.3% Triton X-100 for 2 hours at room temperature. Slices were then immunostained with the appropriate primary antibodies in blocking solution overnight at 4°C on a shaker. Next, the slices underwent three 10 min washes with the blocking buffer and were then incubated with Alexa Fluor 488, 555, or 647-conjugated secondary antibodies for 2 hours at room temperature. After this, three additional washes (15 minutes each) with blocking buffer and a final wash with PBS (10 minutes) were performed. On the penultimate wash 1:1000 Hoechst (Thermo Fisher Scientific, #H3570) was used. The slices were then mounted with fluromount-G (Electron Microscopic Sciences).

### Confocal imaging

Images were captured with a LSM 900 confocal microscopes (Zeiss) using 5x, 10x, 20x or 25x objectives. The same microscope was used for each experiment and identical settings were maintained across all conditions. To image amyloid in the prefrontal cortex, we selected the imaging region based on the mouse brain atlas (Allen Brain Atlas) and located this region based on Hoechst reference. Subsequently, we captured images of a region measuring 200 µm² (3.2055 pixels/µm) through a 20 µm z-stack with 1 µm step sizes. Quantification of images was performed using either FIJI image processing software (v1.54, NIH) or Imaris (v9.1, Bitplane, Zurich, Switzerland). In each experimental scenario, two coronal sections per mouse were utilized from the specified number of animals. The quantification process involved using the averaged values from two to four images per mouse.

### RNA in situ hybridization

We performed RNAscope for fluorescence in situ hybridization according to the manufacturer’s protocol, using PECAM1 to visualize endothelial cells and Nos3 to visualize endothelial nitric oxide synthase expression. Animals were sacrificed immediately after 40Hz stimulation to minimize stimulation-induced degradation of Nos3. Tissue was sectioned at a thickness of 10 µm using a cryostat (Leica) and preserved with desiccants at −80°C until the RNAscope experiment was conducted. All tissue was processed within one week.

### Statistics

Cell count matrices generated from whole brain cFos datasets were compared using custom scripts written in R. All other statistical analyses were performed using GraphPad Prism 10, MATLAB and its Statistics Toolbox (The MathWorks Inc.). The Shapiro-Wilk normality test was used to determine whether the data were normally distributed. For unpaired group comparisons, we used two-tailed, unpaired Student’s t-tests, while for paired group comparisons, we used two-tailed, paired Student’s t-tests. ANOVA, two-way ANOVA, repeated-measures ANOVA, and respective nonparametric tests were used as appropriate. To assess differences between specific groups, post hoc tests were performed. The Tukey test was used to compare between groups when equal variances were assumed. The Sidak test was used to do multiple comparisons in cases where equal variances were assumed. Last, the Games-Howell test was used when equal variances were not assumed. A p-value of <0.05 was considered statistically significant. In figures, significance levels are indicated with black asterisks as follows: \**p* < 0.05, ** *p* < 0.01, *** *p* < 0.0001.

## ACKNOWLEDGMENTS

We are thankful to Dr. Mitchell Murdock, Dr. Liwang Liu, Dr. Chinnakkaruppan Adaikkan, and all Tsai lab members for their helpful comments during preparation of this manuscript. We thank Erica McNamara for mouse care, genotyping and colony maintenance and Ying Zhou for laboratory management. The authors would also like to thank Emily Niederst and Rosalind Firenze for their critical comments. We are also grateful to Kira Poskanzer and Trisha Vaidyanathan for their helpful contributions to a running wheel design. We gratefully acknowledge generous support from the following individuals and organizations: Robert A. and Renee E. Belfer Family Foundation, The Dolby Family, Glenda and Donald Mattes, Kathleen and Miguel Octavio, Amy Wong and Calvin Chin, Lawrence and Debra Hilibrand, Dave Wargo, the Norbert H. Hardner Foundation, The JPB Foundation, and The Picower Institute for Learning and Memory for their generous contributions to our work. Martin Kahn was supported by Swiss National Science Foundation postdoctoral fellowships (<u>P500PM_214140</u> and <u>P2SKP3_199448</u>), Cristina Blanco-Duque was supported by an Alana Down syndrome Center Fellowship. Li-Huei Tsai discloses support for this work from NIH R01AT011460-01 and NIH RO1AG069232.

## AUTHOR CONTRIBUTIONS

NSL, CBD and MCK contributed equally to the project. CBD, MCK and L-HT conceptualized the study. NSL performed whole brain cFos experiments, analyzed data, and interpreted results. MCK and CBD performed fiber photometry experiments, analyzed data, and interpreted results. CBD, MCK and NSL performed *in vivo* electrophysiology experiments and analyzed data. CBD and MCK interpreted results of *in vivo* electrophysiology experiments. NSL, CBD and CYY performed *in vivo* two photon imaging experiments, NSL, CBD and MCK analyzed data and interpreted results. NSL, MCK, CBD, HN, AL, AS performed amyloid immunohistochemistry experiments, analyzed data and interpreted results. NSL and HN performed *in situ* hybridization experiments, analyzed data and interpreted results. AL, RR, AD performed amyloid ELISA experiments, analyzed data and CBD interpreted results. MCK, CBD and AL performed amyloid injection experiments, analyzed data and interpreted results. L-HT provided tools and supervised all aspects of the project. NSL, CBD, MCK, and L-HT wrote the manuscript based on inputs from all authors.

## DATA AVAILABILITY

The authors confirm that the data supporting the findings of this study are available within the article and its supplementary materials. Raw data will be made available upon reasonable request.

