## Supplemental Figures for "The role of cholinergic signaling in multi-sensory gamma stimulation induced perivascular clearance of amyloid"

**Supplementary Figure 1: Whole brain activity mapping. (A)** Bar charts of cFOS+ cell counts in all regions passing uncorrected significance testing comparing GENUS and no stimulation **(B)** representative image of manually segmented Substantia Innominata (SI) labelled with cFOS and ChAT, **(C)** quantification of cFOS/ChAT double positive cells in the SI (top) and total ChAT+ cells (bottom) (unpaired t-test). Scale bar: 1000 *μm*

**Supplementary Figure 2: Changes in vessel diameter during GENUS are blocked by cholinergic antagonists.** Example images showing diameter measurements (white line) in arteries during the glymphatic flux experiments (**B)** change in vessel diameter (mean across mice ± SEM) relative to the first measurement, **(C)** comparison between V and M/S groups at the last time point shown in **(B)** (*$p < 0.05$, paired t-test).

**Supplementary Figure 3: GENUS leads to an increase in endothelial cell NOS3 expression (A)** RNA scope *in situ* hybridization of frontal cortex tissue probed for the endothelial cell marker PECAM1 (pink) and the nitric oxide synthase 3 (NOS3, yellow). Tissue was counterstained with the nuclear marker DAPI. Scale bar: 50 *μm* **(B)** quantification of total number of Nos3 puncta inside of PECAM1-based vessel surfaces (left) and of total total number of PECAM1 punctae (right) (*$p < 0.05$, unpaired t-test).

**Supplementary Figure 4: GENUS leads to increased removal of tissue-injected fluorescent amyolid $\beta_{42}$ (A)** representative images of sequential frontal cortex sections injected with HyLite-amyolid $\beta_{42}$ (red) and collected after 1h of GENUS. Tissue was counterstained with the nuclear dye DAPI. **(B)** quantification of tissue area covered by fluorescent amyloid following GENUS with V injection and **(C)** after injection with cholinergic antagonists (**$p<0.01$, unpaired t-test).

### Supplemental Figure 1

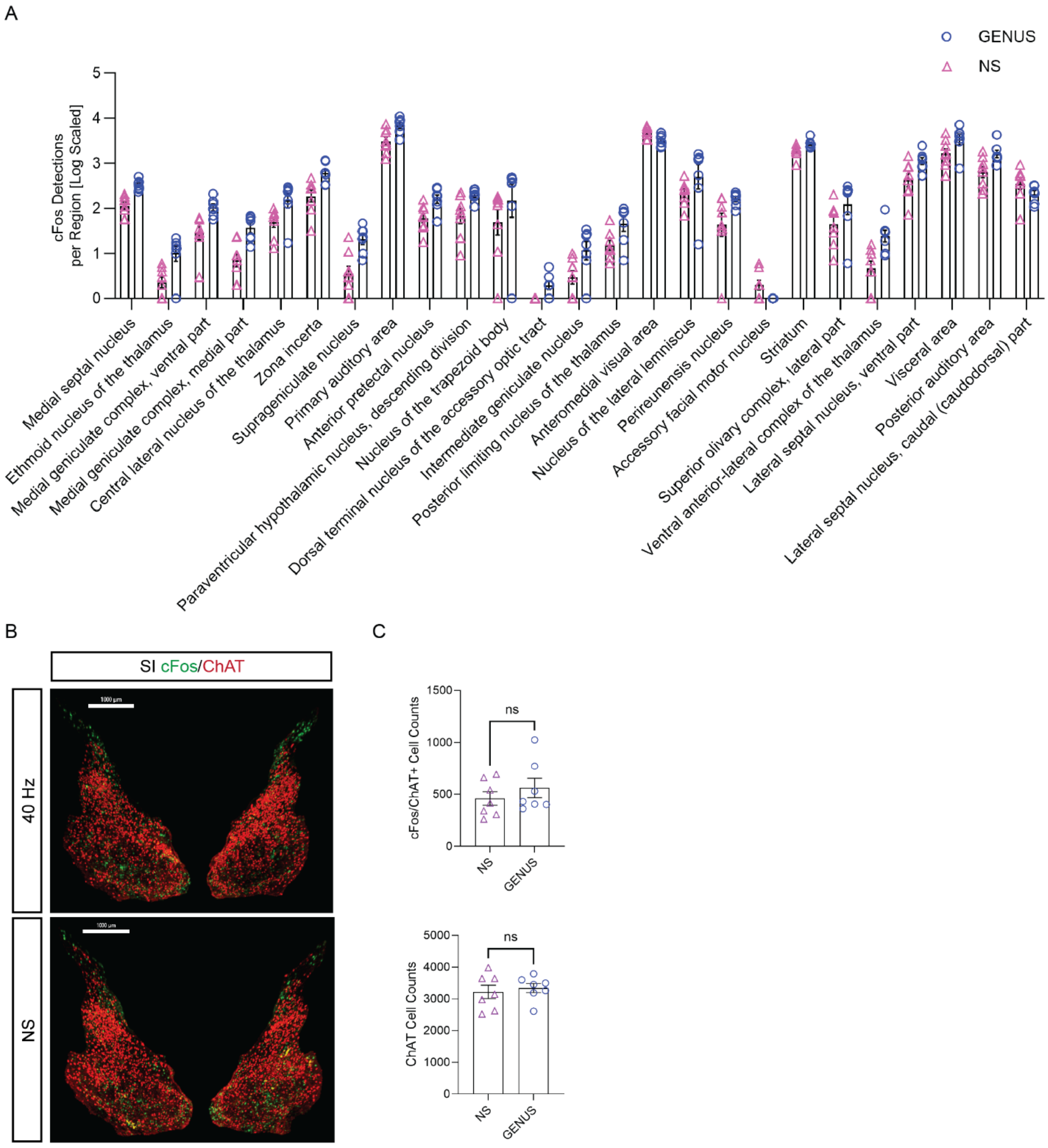

Supplemental Figure 2

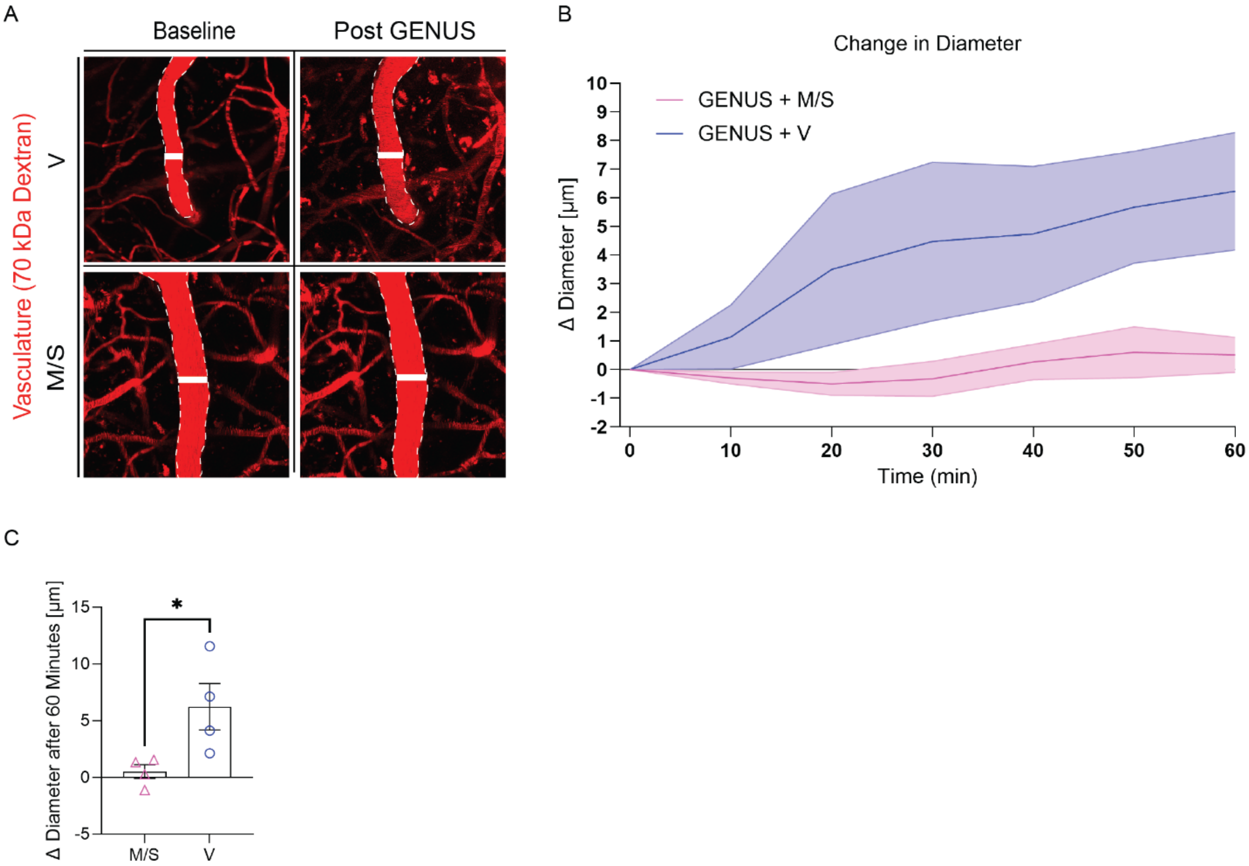

Supplemental Figure 3

A

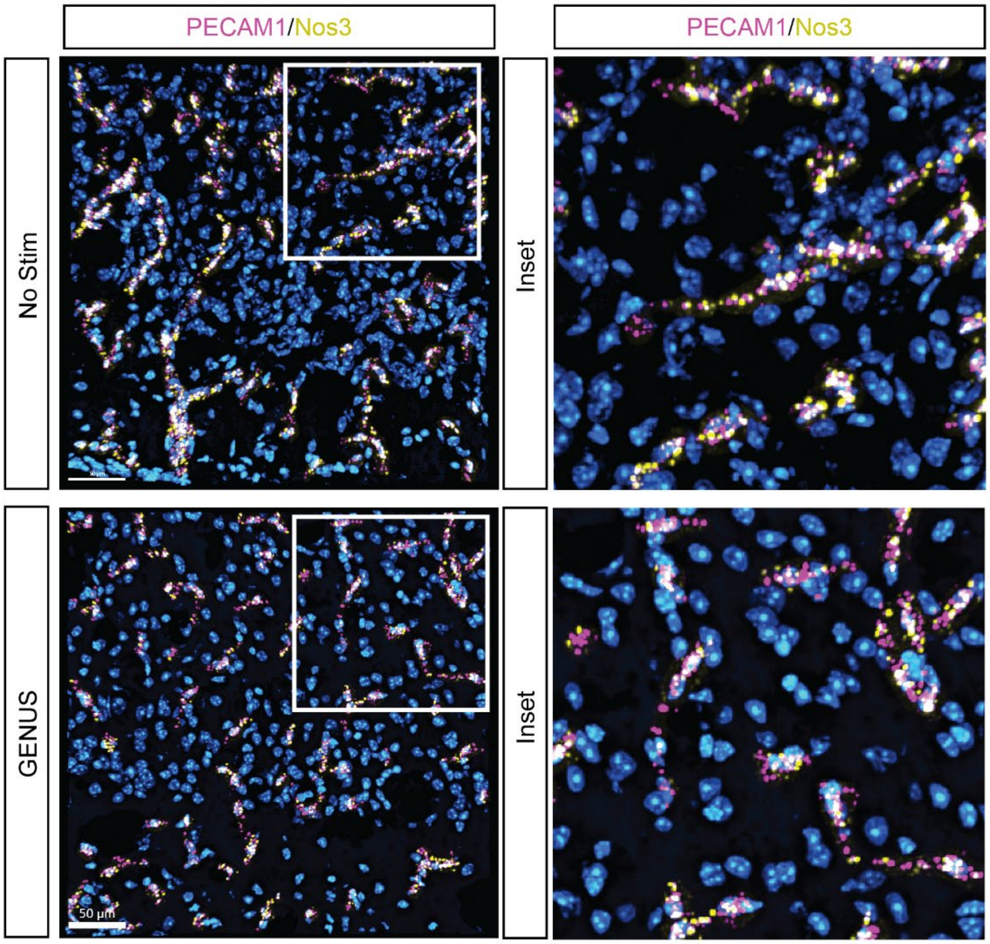


B

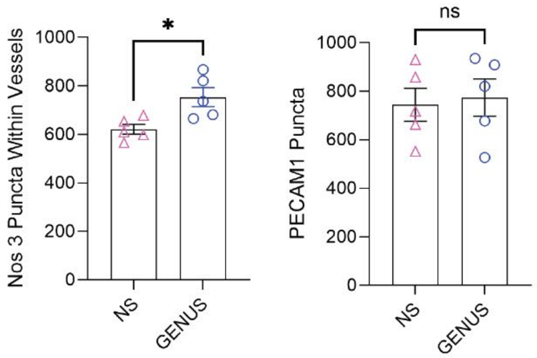

Supplemental Figure 4

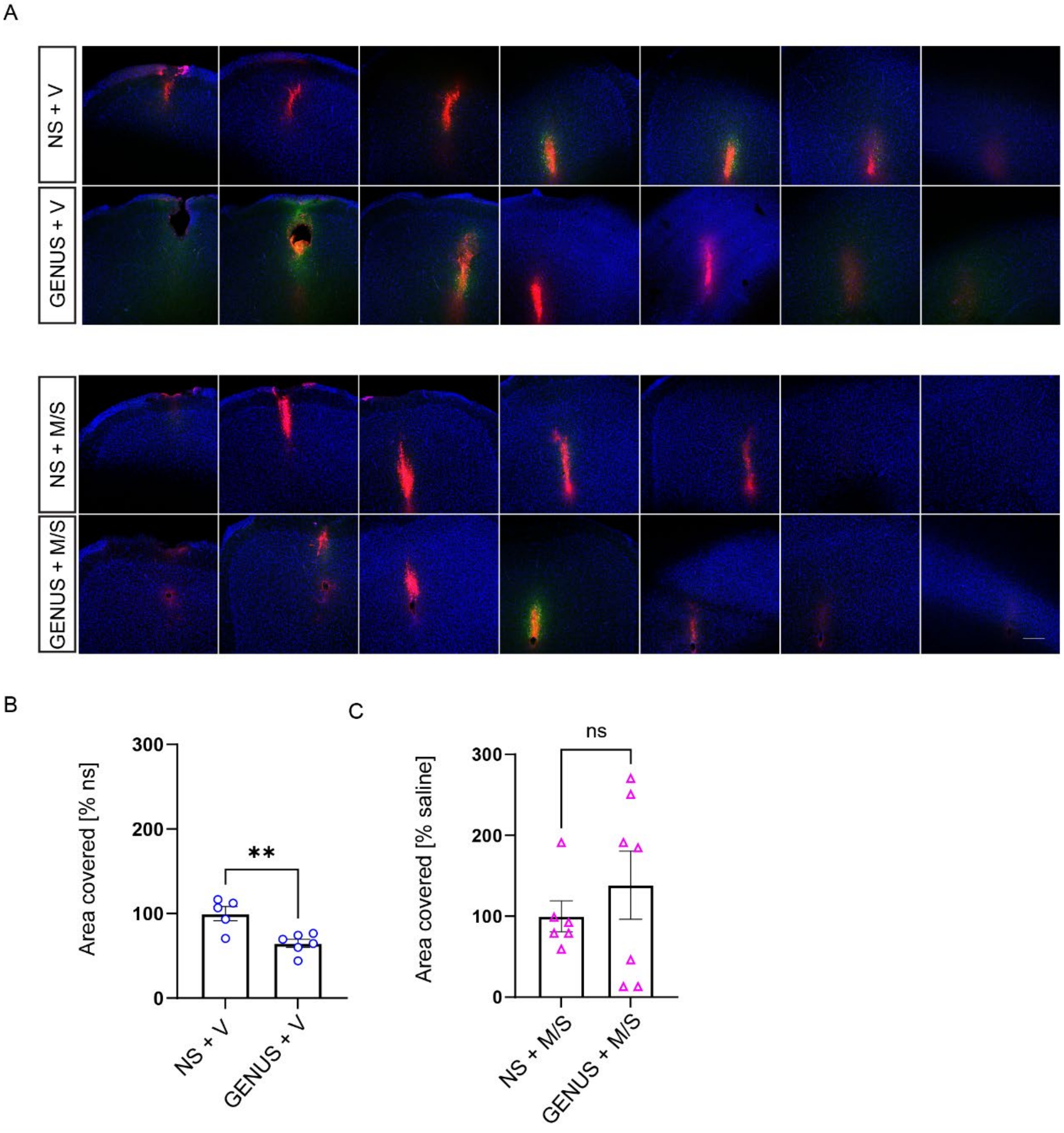
